# Quantifying Uncertainty in the Re-Emergence of Yellow Fever Virus

**DOI:** 10.64898/2026.01.28.702222

**Authors:** Nate Kornetzke, Helen J. Wearing

## Abstract

Emerging infectious diseases are a persistent public health threat that challenge deterministic, mechanistic modeling approaches. Because outbreaks initially start with a low number of infected hosts, their dynamics are highly stochastic, making traditional deterministic methods, e.g. ordinary differential equations, unable to qualitatively or quantitatively capture the transmission dynamics. In place, stochastic models are used, such as Markov chain models, but these models present their own challenges. Often, to infer a quantity of interest with stochastic models, we need to sample the model’s distribution many times over, introducing an additional source of noise to our analysis. This additional noise makes the inference of our quantity of interest more difficult and computationally expensive. Here, we show how novel tools from the field of uncertainty quantification can be used to efficiently separate these two channels of noise, allowing us to make rigorous statistical inferences about processes important for disease emergence. We illustrate these techniques with a model of yellow fever virus spillover in the Americas, a virus that has seen rapid re-emergence amongst multiple hosts and vectors in South America over the last decade. We show that only a handful of parameters’ uncertainties greatly affect variation in cumulative disease incidence. In particular, uncertainty in patch connectivity and non-human primate latency has the greatest impact on the variation of cumulative disease incidence of yellow fever virus across patches.

**Author Contributions:** **Conceptualization:** Nate Kornetzke, Helen J. Wearing. **Methodology:** Nate Kornetzke, Helen J. Wearing. **Formal Analysis:** Nate Kornetzke. **Software:** Nate Kornetzke. **Visualization:** Nate Kornetzke. **Writing, Original Draft Preparation:** Nate Kornetzke. **Writing, Review and Editing:** Helen J. Wearing. **Supervision:** Helen J. Wearing.

**Author Summary:** Emerging infectious diseases are a growing threat to public health. Computational models allow researchers to forecast future disease burdens and to investigate counterfactual scenarios, and often, these models are stochastic, i.e. contain randomness, to capture the qualitative behavior of emergence. A type of statistical analysis known as global sensitivity analysis allows modelers to rigorously analyze how varying the input to a model affects variation in its outputs. This type of analysis helps us infer what mechanisms are important for disease mitigation and control, especially when an infectious disease has a complex ecology consisting of multiple hosts and vectors. Until recently, stochastic models of emerging infectious diseases often proved too computationally expensive to perform a global sensitivity analysis on. Here, we demonstrate how new mathematical tools allow us to streamline this process for complex models of emerging infectious diseases. We analyze the ecology of re-emerging yellow fever in South America, a public health threat occurring in many hosts and vectors across vastly different ecosystems.

## 1 Introduction

Mathematical models have a long, rich history of use in understanding infectious disease ecology. They have been used in forecasting future disease burdens, such as case counts, as well as exploring hypothetical scenarios of mitigation and control [1, 2, 3, 4]. For endemic diseases, deterministic models, e.g. a system of ordinary differential equations, have been successfully utilized in forecasting and mechanistic exploration [1, 2]. However, emergence poses a fundamental challenge to these deterministic methods. Emergence oc-curs when infectious hosts (or vectors) are introduced into a newly susceptible population. The low number of infectious hosts causes stochastic fluctuations, even if the number of new susceptible hosts is large [5]. Indeed, previous modeling work has illustrated the necessity of stochastic models when dealing with disease emergence to avoid erroneous model conclusions [6].

Although stochastic models themselves have a long history of use in infectious disease modeling, they present difficulties in modeling tasks that their deterministic counterparts do not. Almost all dynamic models of infectious disease, both deterministic and stochastic, are analytically intractable, leading us to use computational simulation methods for their analysis instead [2, 5]. For deterministic models, this means simulating a trajectory for a given model input, while for stochastic models, this means simulating a distribution of trajectories for a given model input. In practice, this distribution is inferred via a Monte Carlo method, e.g. Gillespie’s method, simulating many model runs per model input to infer the distribution for that input [7]. For simple models, simulating multiple trajectories for a single model input may only be marginally more expensive. Yet, modelers are increasingly building more complex models to better represent the underlying biology.

Inferring what mechanisms are important in a complex, parameter rich model can be difficult without some notion of which parameters are more influential on model outcome than others. Many model parameters are prescribed biologically meaningful ranges from previous empirical studies, and we can think of these ranges as representing uncertainty in the parameter value. A natural question then arises about how uncertainty in the model parameters affects uncertainty in model outcome or our quantities of interest; this task is known as global sensitivity analysis (GSA) [8]. When the model is additionally stochastic, this can complicate such efforts, as we now have two sources of uncertainty: variation from the stochastic solver itself, i.e. from the Monte Carlo method for inferring the distribution, as well as variation from the parameter ranges. While many methodologies have been established to perform GSA on deterministic models, fewer methodologies have been developed for stochastic models. Recent work in the field of uncertainty quantification has focused on bridging this gap [9, 10, 11, 12]. Here, we showcase the use of some of these new developments and their relevance for infectious disease modelers on a parameter rich model of re-emerging yellow fever virus (YFV) in South America.

YFV is a mosquito-borne virus found throughout the tropics and neotropics that can infect many primate species, including humans. In the 20th century, an effective vaccine was developed and deployed widely across South America, eradicating the virus’s urban transmission cycle. Today, the virus is endemic to non-human primates and forest genera of mosquitoes [13, 14]. Until recently, YFV was largely restricted to heavily forested areas, such as the Amazon. The last decade has seen YFV spread eastward, out of the Amazon, and spillover of YFV from non-human primates to humans has become more frequent, with large outbreaks occurring during 2016-2018 [15, 16]. Genomic evidence revealed that these outbreaks were sylvatic (forest) in origin [17]. Why YFV’s distribution in South America has expanded eastward and spillover has become more frequent is an open question with significant implications for public health and policy. But the ecology of YFV is complex, involving multiple transmission cycles, each with their own primate hosts and vectors [14]. To understand YFV’s re-emergence, we develop a stochastic model of the multiple transmission cycles with multiple hosts and vectors. Because the composition of hosts and vectors across each transmission cycle is unique, we have different parameters associated with each transmission cycle, and consequently, the model is parameter rich. To better understand how variation in these parameters affects variation in YFV incidence across the transmission cycles, we perform a GSA using new tools from uncertainty quantification. Here, we give a brief road map of what follows. First, we describe the ecology of YFV and its emergence, enough to develop our model. From there, we present Sobol indices, a GSA tool used to quantify a model outcome’s sensitivity to uncertainty in model inputs, and how these indices are computed. Then, we describe the variance deconvolution estimator, an estimator that allows one to delineate noise arising from the stochastic solver apart from noise arising from the variation of the parameters themselves, and how we can use it to successfully compute Sobol indices for stochastic models. Subsequently, using parameter ranges ascertained from the literature, we simulate our YFV model and compute its Sobol indices. We use these indices to infer which parameters’ uncertainties have the largest effect on the uncertainty of YFV incidence, and subsequently, we investigate some individual groupings of parameters and how variation across these groupings affects variation of YFV incidence. Finally, we draw some lessons for future investigations of YFV and emerging infectious diseases more generally.

## 2 Yellow Fever Ecology

Originally from Africa, YFV and the competent vector *Aedes aegypti* were brought over to the Americas during colonization and the slave trade [18, 13]. Both became endemic in urban centers, and eventually the virus spilled over into non-human primates (NHPs) living in heavily forested areas, including much of the Amazon. Successful vaccination and vector control campaigns eliminated YFV transmission between humans and A. aegypti in urban areas by the mid-20th century, a transmission cycle known as the urban cycle, but the virus continued to spread unabated among non-human primates (NHPs) in the forest, a transmission cycle known as the sylvatic cycle [13, 14].

Until recently, YFV cases in South America were a result of humans going into forested areas and opportunistically getting bit by an arboreal mosquito that had fed on an infectious NHP. However, the last decade has seen YFV spread eastward, into the Atlantic forest and the Cerrado, with enzootic outbreaks spilling over into urban epicenters like Sao Paulo [17, 15]. In these new areas, spillover has been occurring frequently in rural areas and along city edges, the peri-domestic or peri-urban landscape. Here, various species of primates and mosquitoes co-exist, creating many pathways for pathogen transmission and hence spillover. Intensive land use, like agriculture, has increased the size of these peri-urban landscapes while fragmenting forest and savanna, and it has been postulated in the literature that these areas heighten pathogen spillover [19, 20]. Indeed, while the 2016-2018 outbreaks were largely fueled by arboreal species of mosquitoes from the genera *Haemagogus* and *Sabethes*, there are ample ecological conditions for a transmission cycle to occur between the primate hosts and mosquito vectors in the peri-urban landscape, the peri-urban cycle. Already, there is empirical evidence of YFV circulation in a key peri-urban vector, the highly opportunistic feeder *A. albopictus*, and monitoring of forest edges has revealed a significant level of YFV incidence in NHPs [21, 22, 23]. The public health threat posed by emerging YFV highlights the need for understanding these ecological conditions and how they could lead to YFV spillover. Indeed, we examine the hypothesis that the peri-urban cycle acts as a highway between the two cycles, amplifying the probability of spillover, with the different vector ecology across the cycles also playing a prominent role in spillover incidence [16, 15, 14, 20, 19].

To investigate this hypothesis, we turn to mathematical modeling. The ecology of YFV transmission across these three cycles — the sylvatic, the urban, and the periurban — is complex. Each cycle involves different host and vector species with different life histories, and the cycles are loosely connected to one another. Additionally, spillover typically occurs with low numbers of infected hosts, and thus, stochasticity plays a prominent role in the emergence of YFV. These factors make elucidating the mechanics of YFV spillover difficult through empirical studies alone, and mathematical epidemiology contains a rich literature on approaching infectious disease systems marked by nonlinearity and stochasticity [2].

The last two decades has seen a growing body of modeling literature on the sylvatic cycle of mosquito-borne viruses, including YFV. In 2012, Althouse et al. [24] used a stochastic, multi-host SEIR-style compartmental model to demonstrate that the sylvatic cycle of dengue in Senegal likely relies on stochastic introductions. Later, this model was expanded upon to investigate the probability of sylvatic Zika taking off in the Americas [25]. However, both of these models lacked environmental or climatic factors and their effects on YFV transmission. In 2019, a mechanistic probability model was devised by Childs et al. [26] to predict YFV spillover incidence across Brazil as a function of climate and land use, highlighting the importance of these variables in assessing YFV threat. Motivated by these works, our model explores the peri-urban cycle acts as a conduit between the urban and sylvatic cycle.

## 3 The Model

We construct a continuous-time Markov chain model (CTMC) to model spillover from the sylvatic cycle into the peri-urban and urban landscapes.

Each cycle contains a vector population and at least one host population, and each population is split into compartments based on their infection status, similar to the susceptible-exposed-infectious-recovered paradigm [1, 2]. The sylvatic cycle contains a non-human primate population and a mosquito population, while the urban cycle contains a human population and a mosquito population. The peri-urban cycle contains both a human population and a NHP population as its hosts, but it contains one mosquito population that feeds on both. The peri-urban cycle is connected to the urban cycle via human movement and to the sylvatic cycle via NHP movement, as illustrated in figure 1.

**Figure 1:**
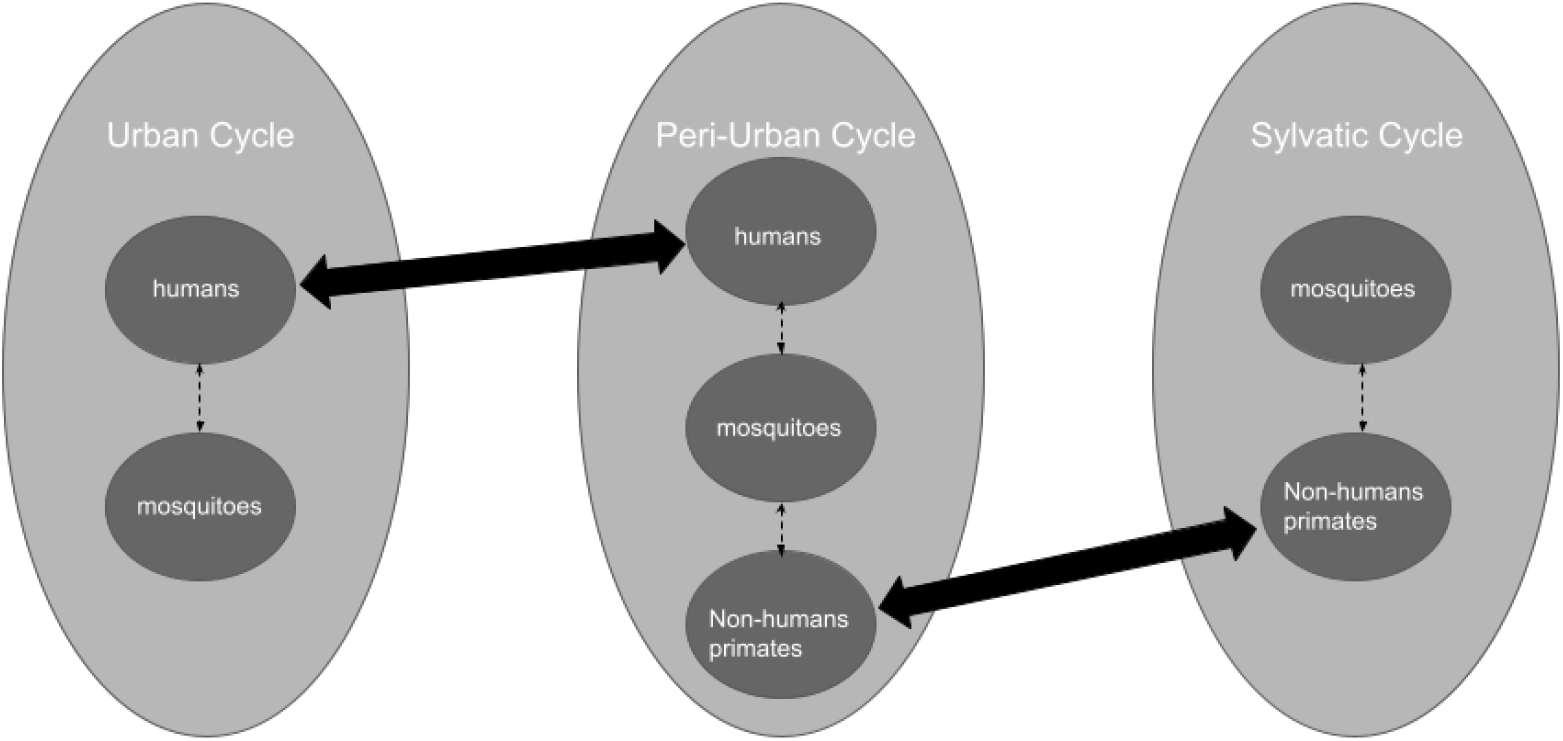
Conceptual model of the three transmission cycles, an extension of the SEIR-SEI model for mosquito-borne pathogens to three patches. Here, each patch contains at least one host population, either humans or NHP, and one mosquito population. Each of these populations are divided into compartments based on infection status, as is typically done in a SEIR-SEI model. The patches are connected by movement between hosts (solid arrows). Connectivity between the urban and peri-urban cycles results from human movement, while the connectivity between the peri-urban and sylvatic cycle comes from NHP movement. Each host and vector population within a cycle are parameterized to key species for that transmission cycle (dashed arrows), e.g. the host population in the urban cycle is parameterized to humans and the mosquito population is parameterized to *A. aegypti*. We do not consider human movement from the urban cycle to the sylvatic cycle because we are primarily interested in the role that the peri-urban cycle plays as a bridge between the sylvatic and the urban cycles.

## 3.1 State variables

Each cycle’s host and vector populations are denoted by 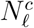, where *c* = *u, p, s* for urban, peri-urban, and sylvatic, and *𝓁* = *h, p, m* for human, non-human primate, and mosquito, respectively. Throughout this paper, superscripts will refer to which cycle a state variable or parameter belongs to, while subscripts refer to which species of host or vector. We illustrate how these populations are further structured by discussing the urban cycle host and vector populations, but we note that each population in each cycle follows the same structure unless stated otherwise.

The urban human population, 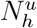, is divided into four compartments: susceptible, exposed, infectious, recovered. The susceptibles, 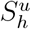, are those who have not been exposed or infected to YFV. The exposed, 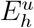, are those who have been infected by an infectious mosquito but are not yet infectious; that is, they are in the latent period. The infectious, 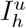, are those who are infectious following exposure. The recovered, 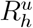, are those convalescing or fully recovered. YFV infection provides lifelong immunity, and thus the transmission cycle ends in recovery [14]. The urban mosquito population, 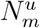, is divided into three compartments: 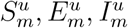. Adult female mosquitoes are recruited susceptible, 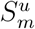, and become exposed, 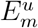, after biting an infectious primate. Infectious mosquitoes, 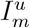, stay infectious for the rest of their life, as mosquito lifespans are not long enough to clear viral infection [14].

### 3.2 Equations

Movement between the compartments occurs with probabilities which functionally depend on fixed parameters and the state variables themselves. We first motivate the probabilities within one transmission cycle by constructing the urban cycle. We follow a similar construction as the Ross-Macdonald model [27]. The full set of events and their probabilities are found in the SI.

Susceptible humans, 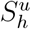, become infected after being bit by an infectious mosquito, moving them to the exposed class,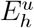. This rate relies on how frequent a mosquito bites primates, given by a biting rate *b*^*u*^. Now, 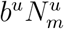 gives the total number of mosquito bites, and 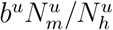 gives the total number of mosquito bites per person. Thus, 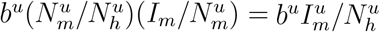 gives the total number of infectious mosquito bites per person. YFV transmission does not always occur when an infectious mosquito bites a susceptible primate; instead, it transmits with a probability, given by *β*_*mh*_. Of course, transmission depends on how many susceptible hosts there are, 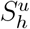.

Once successful transmission of YFV from an infectious mosquito to a susceptible human occurs, the primate is not infectious right away. Indeed, YFV has a several day latent period before the primate host can infect a susceptible mosquito that bites it. Thus, individuals leave the exposed class and become infectious at a rate 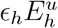, where *ϵ*_*h*_ is inversely proportional to the latency period. Similarly, the infectious period of YFV can last up to a week, but after that, an individual is convalescing, and so infectious individuals move to 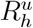 at a probability 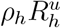, where *ρ*_*h*_ is inversely proportional to the infectious period. We do not consider background birth or death of humans in either the urban or the peri-urban cycle because we simulate our model at a short enough time scale where those occur at negligible rates relative to the time scale. We do not consider diseaseinduced mortality for humans because disease incidence of humans occurs independently of whether someone in the recovered class, i.e. someone convalescing, succumbs to diseaseinduced mortality or not.

The transmission dynamics for the adult female mosquitoes 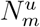, see figure 2, are similar to those of the host. A susceptible mosquito, 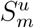, becomes infected when it bites an infectious primate, in the urban case a human, moving it to 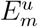. The rate of this event depends on the amount of bites a mosquito can take from a human, 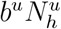, but this is per human, so we get 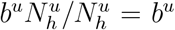. Like the transmission term previously discussed, this event relies on a rate that transmission actually occurs, *β*_*hm*_, as well as the proportion of infectious humans 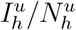 and the number of susceptible mosquitoes 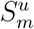. We give the event and its rate in table 1.

**Table 1:** A CTMC is fully described by its events and the rates of those events; here, we give a subset of the events and rates that describe our CTMC. Recall that in a CTMC, the state variables are integers, so the events describe changes in the integer values of the state variables. For example, 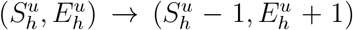 tells us that 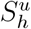 decreases by 1 and 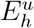 increases by 1, and all the other state variables in the model stay the same. The rates are functions of the state variables and the model parameters, and they are used to compute the probability of each event occurring. The event 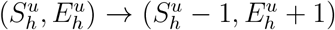 occurs at the rate 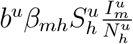, where *b*^*u*^ and *β*_*mh*_ are fixed parameters while 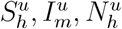 are the values of those state variables at the time the event occurs. Since our model has the Markov property, we assume that the time to the next event is exponentially distributed.

| Biological Description | State Variable Change | Rate |
| --- | --- | --- |
| **Host Transmission** |  |  |
| Susceptible host infected by infectious mosquito, entering a latency period | $(S_h^u, E_h^u) \rightarrow (S_h^u - 1, E_h^u + 1)$ | $b^u \beta_{mh} S_h^u \frac{I_m^u}{N_h^u}$ |
| **Mosquito Transmission** |  |  |
| Susceptible mosquito infected by an infectious host, entering a latency period | $(S_m^u, E_m^u) \rightarrow (S_m^u - 1, E_m^u + 1)$ | $b^u \beta_{hm} S_m^u \frac{I_h^u}{N_h^u}$ |
| **Mosquito Demographics** |  |  |
| Mosquito recruitment | $(S_m^u, N_m^u) \rightarrow (S_m^u + 1, N_m^u + 1)$ | $\Theta^u N_m^u$ |
| Mosquito death | $(S_m^u, N_m^u) \rightarrow (S_m^u - 1, N_m^u - 1)$ | $\frac{\Theta^u}{K^u} N_m^u S_m^u$ |
| **NHP Demographics** |  |  |
| NHP birth | $(S_p^s, N_p^s) \rightarrow (S_p^s + 1, N_p^s + 1)$ | $\nu_p N_p^s$ |
| NHP background mortality | $(S_p^s, N_p^s) \rightarrow (S_p^s - 1, N_p^s - 1)$ | $\mu_p S_p^s$ |
| NHP disease-induced mortality | $(R_p^s, N_p^s) \rightarrow (R_p^s - 1, N_p^s - 1)$ | $\eta_p R_p^s$ |
| **Travel** |  |  |
| Susceptible host movement from the urban cycle to the peri-urban cycle | $(S_h^u, N_h^u, S_h^p, N_h^p) \rightarrow (S_h^u - 1, N_h^u - 1, S_h^p + 1, N_h^p + 1)$ | $\tau_u S_h^u$ |

**Figure 2:**
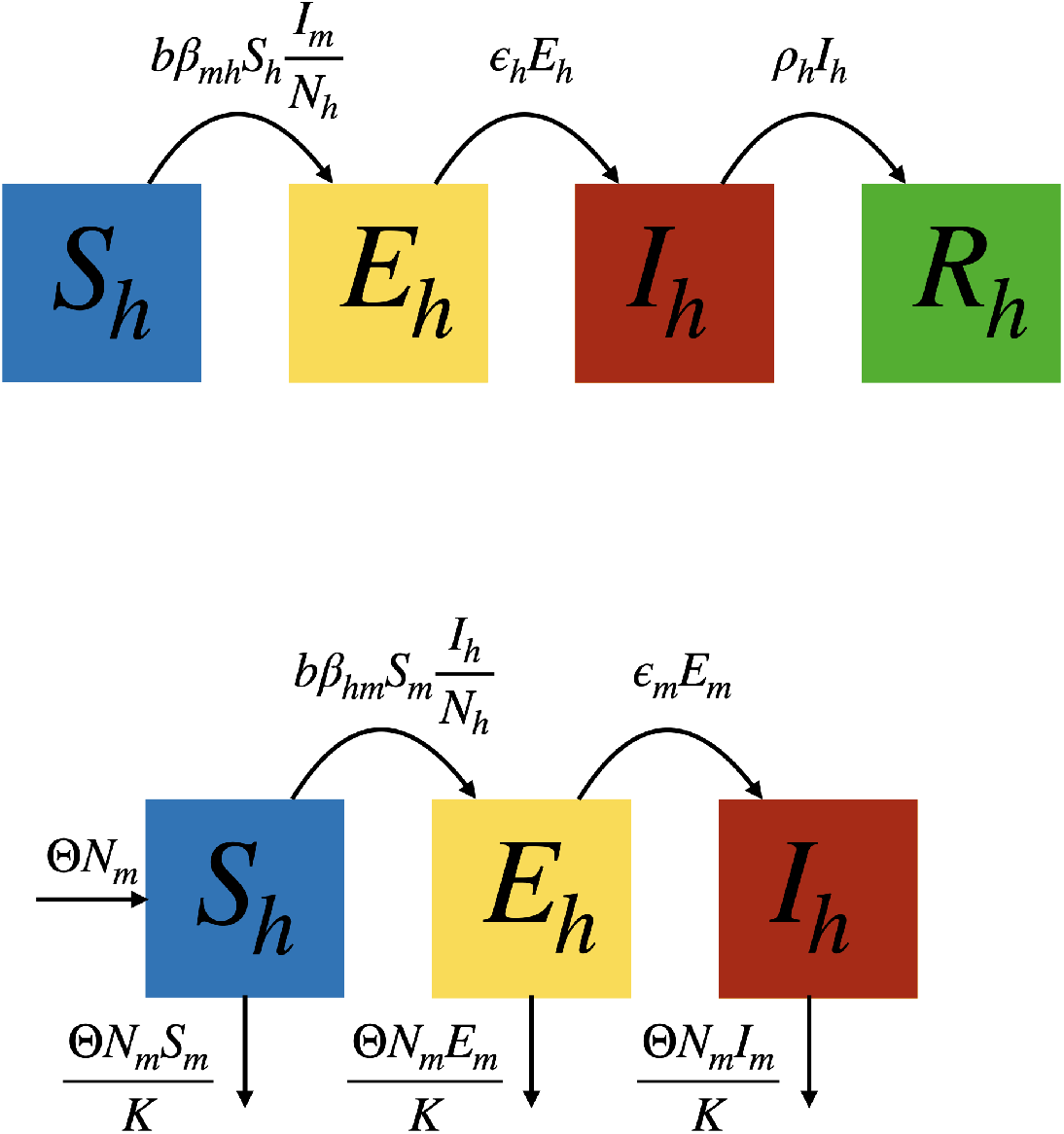
Each host or vector population is split into compartments with respect to infection status — susceptible, exposed, infectious, and recovered — and transitions between compartments occur at rates dependent on timeconstant parameters and the state variables. Here, we assume frequency-dependent infection, an assumption that matches the observed disease dynamics of YFV and other mosquito-borne viruses. For NHP, we include both background birth and deaths, as well as disease-induced mortality because NHP demographic processes occur at time scales smaller than the length of simulation, and NHP deaths from YFV can be catastrophic, affecting how many new susceptibles are recruited [28]. For mosquitoes, we exclude a recovered compartment, as they do not live long enough to recover from infection, remaining infectious for the remainder of their life. Births and deaths are density dependent. The carrying capacity K is set to be twice the host size, assuming 2 mosquitoes per person.

After infection, the mosquito moves from 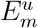 to 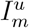 at a rate 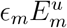, where *ϵ*_*m*_ is inversely proportional to the latency period. Mosquitoes in 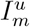 do not recover. However, mosquitoes in any of the three compartments can die, and new adult female mosquitoes can be recruited into 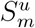. We do not consider the full life history of the mosquito, i.e. the juvenile stages, for model simplicity. The birth and death probabilities occur with a probabilistic analogue to the logistic growth equation, given in table 1.

The probabilities for the transmission cycle between NHPs and mosquitoes in the sylvatic cycle are similar to the probabilities described above. However, unlike humans in the urban or peri-urban cycle, we include births and deaths for NHPs. Smaller NHPs have much shorter lifespans than humans, and we consider it important relative to the time-scale of our model. We also include death by disease for NHPs, as neotropic NHPs have shown considerable levelsmodel diagram of mortality to YFV, with mortality rates ranging from 1% to over 60% depending on the species of NHPs [28].

The human transmission events in the peri-urban cycle are almost the same as those in the urban-cycle, but the transmission term is modified. The biting rate *b*^*p*^ occurs over all possible hosts, 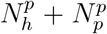, not just humans, 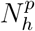. This gives us the transmission term 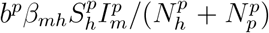. This change is also seen in the transmission term for the peri-urban NHPs and the peri-urban mosquitoes.

The three cycles are linked together by primate movement. Humans move between the urban and peri-urban cycle with the rates *τ*_*u*_ and *τ*_*p*_, while NHPs move between the peri-urban and sylvatic cycle with the rates *φ*_*p*_ and *φ*_*s*_. This movement occurs with a constant rate proportional to compartment size, with an example of such an event and its rate in table 1. Movement between infectious compartments does not occur, as we assume that sick primates do not move enough for our model to consider.

### 3.3 Quantities of Interest and Hypotheses

We are investigating the emergence and spillover of YFV into the hosts of the periurban and urban transmission cycles, so we consider cumulative disease incidence of the hosts in each cycle as our quantities of interests (QoI). This QoI tells us whether YFV is occurring in a particular transmission cycle or not for a given set of parameters. As previous literature has highlighted the importance of movement across ecosystems, we hypothesize that variation in the travel parameters will contribute the most to variation of our QoIs, and we also hypothesize that variation in certain biotic parameters, particularly the primate birth rate, will also contribute heavily to the variation of QoIs [20].

## 4 Methods

### 4.1 Model Simulation, Initial conditions, and Parameter Values

Investigating how uncertainty of the parameter values affects the uncertainty of the QoIs requires being able to compute the distribution of the CTMC for a given parameter value, which is done by solving the Kolmogorov forward equations of the CTMC (see SI for more details). Solving these equations is often analytically and numerically intractable. Instead, Monte Carlo methods, such as the stochastic simulation algorithm (SSA), have been developed to infer the distribution by simulating trajectories whose events adhere to the underlying distribution. Approximation methods, known as tau-leaping, have subsequently been developed to reduce the computational cost associated with the original exact method. However, some tau-leaping methods are known to over count events, producing negative state variables when the state variables get close to 0 [29]. Many methods have been developed to avoid this dramatic over counting, including the binomial tau-leaping method. Since our simulation starts with an initial condition where many state variables are close to 0, we are essentially prone to this issue if we use the traditional tau-leaping approach. So, we use the binomial tau-leaping method [30].

We choose an initial state where there is no disease in the urban or peri-urban cycles, and 1 infected NHP in the sylvatic cycle. The values chosen for each cycle’s host populations were based on population densities of 1km^2^ found in the literature [31], [32], [33], [34], [35]. Parameter ranges, discussed in more detail below, were found in the literature as well. We then simulate the model over 12 years. This time scale was chosen because it is roughly the average lifespan of a larger non-human primate, and previous modeling on arboviruses in non-human primates has implicated the importance of primate lifespan in disease incidence [25].

### 4.2 Global Sensitivity Analysis

GSA is a statistical analysis where we compute the variance of a QoI given variance across all the model inputs [8]. If the variance of an input parameter is high but the QoI’s variance is low, then we say that input parameter has low or trivial effect. Conversely, if the variance of an input parameter is high and the QoI’s variance is high, then we say that parameter has high or non-trivial effect. We can quantify these assertions by using Sobol indices [36].

#### 4.2.1 Sobol Indices

Sobol indices quantify the variance of a model QoI attributable to a particular model parameter or set of model parameters. It has been demonstrated that the total variance of a square-integrable function of *n* inputs can be decomposed into a sum of variances, giving us the following decomposition [36]:

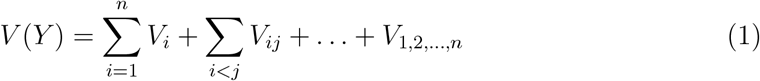

Where

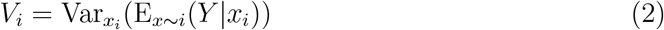

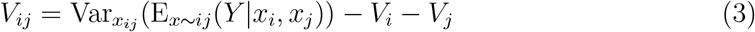

and so on, noting that the notation *x* ∼ *i* means for all parameters except the *i*th parameter. This decomposition allows us to then compute the percentage of variance attributable to a particular parameter, the first order Sobol indices, as well as the percentage of variance attributable to interactions of parameters, the higher order Sobol indices.

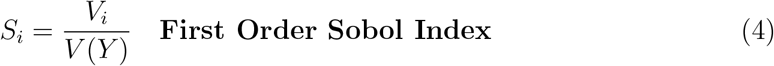

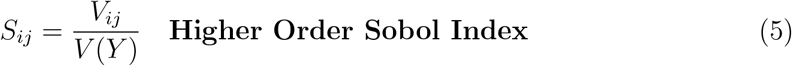

In practice, we have 2^*n*^ − 1 possible Sobol indices to compute, and since computation of the Sobol indices, discussed shortly, can be time consuming, we typically do not compute every Sobol index of a model. Instead, we compute the first order indices along with what are called the total order indices, which tell us how much variance in the model is due to the sum of first order and higher order effects.

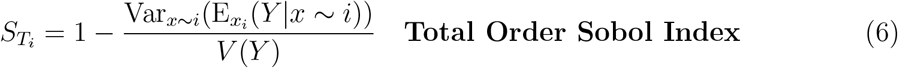

Together, the first order and total order Sobol indices inform of us of how influential a model parameter is on a QoI without having to compute every higher order index involving that parameter. To compute these indices for a model QoI, a Monte Carlo procedure is taken. First, a region of parameter space is sampled, the model is run on each parameter sample to compute the QoI, and then equations (2) and (6) are computed using the standard estimators for expectations and variances. However, notice that we have an expectation inside a variance in both equations. This causes a double loop in our Monte Carlo procedure. That is, if we have *N*_0_ samples of parameter space, then we must run our model 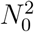 times to compute quantities on the right-hand sides of equations (2), (6). This quickly becomes prohibitive for more complex models that take time to simulate.

Fortunately, Saltelli devised an efficient sampling scheme that reduces the model runs required to compute the first and total order indices from 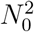 down to *N*_0_(*n* + 2) [8]. Instead of sampling *N*_0_ *× n* samples from parameter space, we sample *N*_0_ *×* 2*n* samples, splitting them into matrices *A* and *B*. Then, we form *n* matrices *C*_*i*_ by replacing the *i*th column of *B* with the *i*th column of A. We run our model on the rows of each matrix, denoting the output by 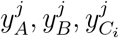 where *j* = 1, …, *N*_0_. Then, we can compute the Sobol indices using the equations below.

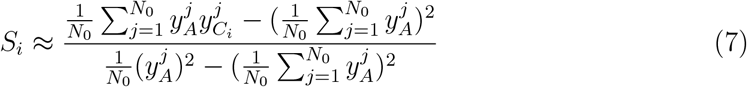

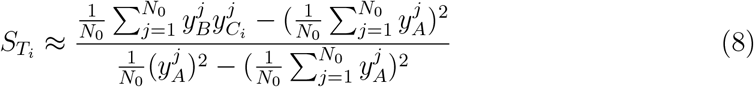

#### 4.2.2 Stochastic Solvers and Variance Deconvolution

While Saltelli’s sampling scheme reduces the number of model runs from 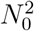 to *N*_0_(*n* + 2), we are faced with another computational burden when we use a stochastic model. CTMCs are rarely analytically tractable, so instead, we compute the QoIs using a Monte Carlo simulation algorithm, such as the SSA, where we simulate a trajectory that belongs to our model’s probability distribution [29]. Since it is a Monte Carlo procedure, we introduce an additional source of variance by running the simulation model many times to resolve the QoIs. A naive way to reduce the solver variance is to over-resolve the model by making the number of model runs *k* large. This makes the number of required model runs using equations (7), (8) to *k×N*_0_(*n*− 2), and so if *k* is moderately large, this becomes infeasible for any mildly complex stochastic model.

Fortunately, the variance deconvolution estimator allows one to avoid over-resolving

the stochastic solver by explicitly accounting for the variance produced by the stochastic solver. It does this by computing the “polluted” variance estimate 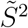 of our model, i.e. the usual estimator of variance, and the variance of the solver, taking the difference of the two to get a “parametric” variance *S*^2^, which we can think of as the “true” model variance [11]. This estimator has been shown to be unbiased while reduandersoncing the number of model runs for a stochastic solver necessary to compute the model variance. Here, we review how it is used by Clements and others to compute Sobol indices [12].

If *Q* is the QoI of our CTMC, then we have a stochastic solver *f* (*x*; *j*), where *x* is the model parameters and *j* is the *j*th run of *f*, which produces our estimate of the *Q* using *k* runs:

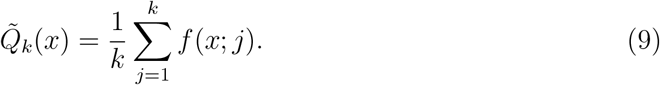

We can now compute the variance of the solver:

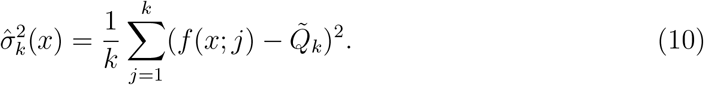

Now, let us sample our parameter space through Saltelli’s method with the matrices *A, B, C*_*i*_. When we run our model on each sample (one of the rows of the matrices), we run the model *k* times and use equation (9) to have the model outputs 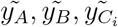. We can likewise compute the solver variance, equation (10), for each model run of matrix *A*, giving us a column vector of solver variances:

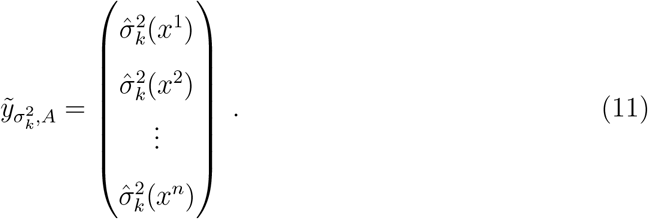

We then estimate our polluted variance:

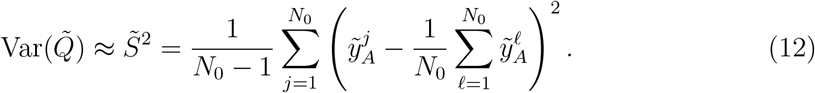

Together with the solver variances, this gives us our parametric variance estimate:

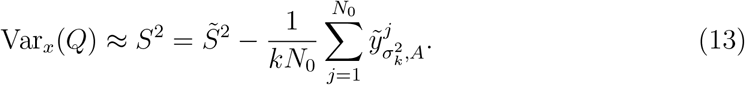

Finally, we can use equations (12), (13) to estimate first and total order indices similarly to how we estimated them with equations (7), (8), except explicitly accounting for solver variance. We get following the estimators:

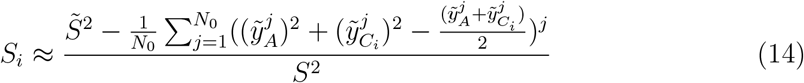

And

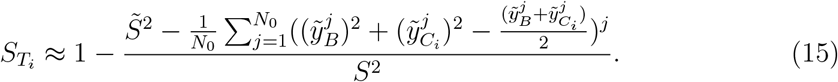

These estimators allow us to feasibly and rigorously compute the first order and total order Sobol indices of a stochastic model. We now do this on our model of YFV across the three transmission cycles, with our QoIs being the mean cumulative disease incidences specified above.

#### 4.2.3 GSA of the three patch YFV model

To perform a GSA on our YFV model, we selected candidate species to represent each cycle’s hosts and vectors, serving as a baseline parametrization. The urban host and vector are represented by humans and *A. aegypti* respectively. One of the peri-urban hosts is humans, while its vector is *A. albopictus*. Since a variety of NHP are infected with YFV in the Americas, we focused on the two most affected genera, the marmosets, *Callithrix*, and the howler monkeys, *Aloutta*. Statistics on species from these genera were used for the NHP host in both the peri-urban cycle and the sylvatic cycle. Similarly, many species of mosquitoes are involved in YFV transmission in forested areas, but we focused on two genera most implicated in YFV’s sylvatic cycle, *Sabathes* and *Haemagogus*, to represent the vector of the sylvatic cycle.

We performed a literature review to determine biologically meaningful parameter value ranges, with each parameter being determined as an annual rate or probability. This is provided in table 2. We subsequently sampled the parameter space uniformly using a quasi-Monte Carlo sampling scheme.

**Table 2:** Parameters ranges used to compute the Sobol indices determined from the literature. Point estimates were turned into ranges by assuming some level of variation around the point estimate. We let parameters with more uncertainty around them have wider ranges. Some of the parameters could not be determined from the literature alone. The sylvatic mosquito reproduction rate was found by widening the ranges of the urban and peri-urban mosquitoes. The transmission probability of NHP to sylvatic mosquito was found by taking the higher ranges of that found in the experimental data for the urban and peri-urban mosquitoes. We postulate there’s a higher transmission probability in the sylvatic cycle because of YFV’s endemic status there. The transmission probabilities of mosquitoes to humans were not found either. YFV vaccination is high in much of South America, and outbreaks are still rare [14]. This used the same ranges determined for NHP. Finally, we note that the travel rates were determined by numerical experiments, picking a wide range that allowed for stochastic die-off as well as epidemics.

| Parameter | Interval | Biological Interpretation | Citation |
| --- | --- | --- | --- |
| $\mu_p$ | [0.05, 0.083] | NHP death | [37], [38] |
| $\nu_p$ | [0.1, 0.25] | NHP birth | [39], [40] |
| $\Theta_u$ | [0.15, 0.45] | urban mosquito reproduction rate | [41] |
| $\Theta_p$ | [0.11, 0.35] | peri-urban mosquito reproduction rate | [42] |
| $\Theta_s$ | [0.17, 0.47] | sylvatic mosquito reproduction rate | N/A |
| $b_u$ | [21024, 46428] | urban mosquito biting rate | [43] |
| $b_p$ | [5383.75, 17976.25] | peri-urban mosquito biting rate | [44] |
| $b_s$ | [11315, 30112.5] | sylvatic mosquito biting rate | [45] |
| $\beta_{hm}^u$ | [0.08, 0.16] | urban human to mosquito transmission probability | [46] |
| $\beta_{hm}^p$ | [0.02, 0.2] | peri-urban human to mosquito transmission probability | [46] |
| $\beta_{pm}^p$ | [0.02, 0.2] | peri-urban NHP to mosquito transmission probability | [46] |
| $\beta_{pm}^s$ | [0.2, 0.35] | sylvatic NHP to mosquito transmission probability | N/A |
| $\beta_{mh}^u$ | [0.25, 0.75] | urban mosquito to human transmission probability | N/A |
| $\beta_{mh}^p$ | [0.25, 0.75] | peri-urban mosquito to human transmission | N/A |
| $\beta_{mp}^p$ | [0.25, 0.75] | peri-urban mosquito to NHP transmission probability | [28] |
| $\beta_{mp}^s$ | [0.25, 0.75] | sylvatic mosquito to NHP transmission probability | [28] |
| $\epsilon_h$ | [0.008, 0.016] | human latency rate | [47] |
| $\epsilon_p$ | [0.0014, 0.012] | NHP latency rate | [48] |
| $\epsilon_m^u$ | [0.019, 0.04] | urban mosquito latency rate | [46] |
| $\epsilon_m^p$ | [0.019, 0.04] | peri-urban mosquito latency rate | [46] |
| $\epsilon_m^s$ | [0.019, 0.04] | sylvatic mosquito latency rate | N/A |
| $\rho_h$ | [0.000391, 0.000913] | human recovery rate | [49] |
| $\rho_p$ | [0.000391, 0.000913] | NHP recovery rate | [48] |
| $\eta_p$ | [0.01, 0.62] | NHP disease death | [28] |
| $\tau_u$ | [0, 1] | human urban to peri-urban travel rate | N/A |
| $\tau_p$ | [0, 1] | human peri-urban to urban travel rate | N/A |
| $\varphi_p$ | [0, 1] | NHP peri-urban to sylvatic travel rate | N/A |
| $\varphi_s$ | [0, 1] | NHP sylvatic to peri-urban travel rate | N/A |

Subsequently, we ran the model on each parameter sample 30 times using binomial tau leaping [30]. We then computed the first order and total order Sobol indices using the estimators (14), (15). Recall that Sobol indices can be interpreted as a percentage, since they are normalized by the model outcome’s total variance. We sampled the parameter space until we had convergence of the Sobol indices up to two decimals. Convergence was determined by increasing the number of parameter samples until the first two decimals remained identical for each estimate. Pre-processing and post-processing of data was performed in the programming language R with RStudio, and the stochastic simulations were done in C++.

## 5 Results

### 5.1 Sobol Indices

The majority of the model parameters have zero first order and total order Sobol indices for our QoIs, indicating that the ranges of uncertainty associated with these parameters do not produce significant uncertainty in cumulative disease incidence across hosts and cycles. This is not to say the majority of the parameters themselves are not important to cumulative disease incidence, but rather that these parameters’ ranges are sufficiently narrow to produce negligible variation in cumulative disease incidence. Six parameters have non-trivial Sobol indices, either first order or total order, and these parameters and their indices are shown in figure 3. The parameters *µ*_*p*_, the background NHP death rate, and *ν*_*p*_, the NHP birth rate, have identical first order and total order Sobol indices for cumulative disease incidence in the sylvatic cycle, indicating that these parameters have entirely first order effects. In panel (d) of figure 3, we see that *ν*_*p*_ accounts for 99% of the variation of this QoI, with the remaining 1% attributed to *µ*_*p*_. Thus, the main driver of variation in cumulative disease incidence in the sylvatic cycle is the direct variation of *ν*_*p*_. This corroborates previous modeling work on the sylvatic cycle that implicated the NHP birth rate as a large driver of sylvatic epidemics [25].

**Figure 3:**
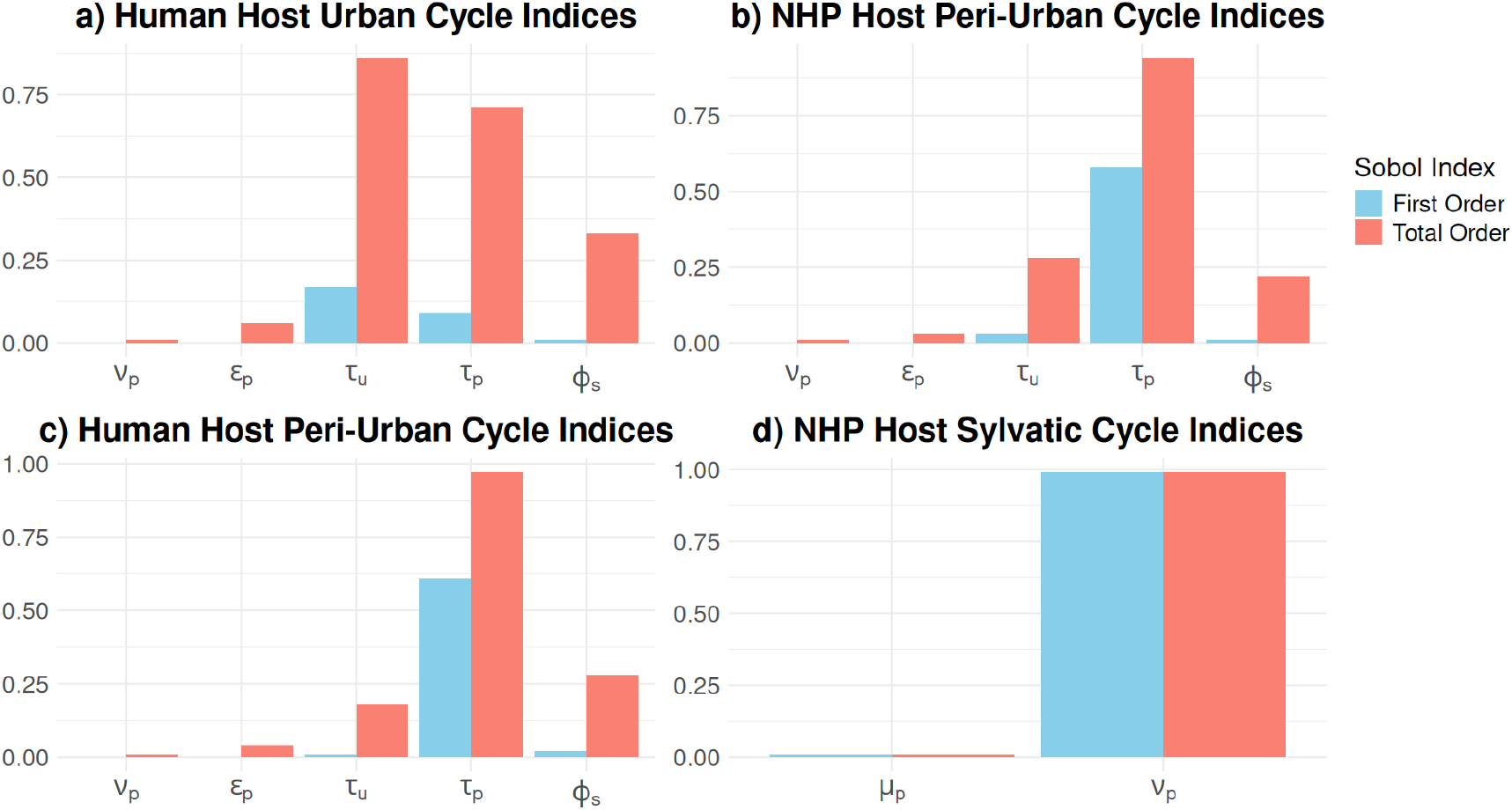
The first order and total order Sobol indices for disease incidence across hosts and cycles demonstrate that variation in only a handful of parameters has significant effect. Here, we show the parameters with non-zero first order or total order indices. Any parameter not shown has zero first order and total order Sobol indices. Disease incidence in the sylvatic cycle has just two significant parameters, the death rate *µ*_*p*_ and the birth rate *ν*_*p*_. We see that the first order and total order indices are identical for both parameters, meaning these parameters both only have first order effect on disease incidence of NHP in the sylvatic cycle. In the other cycles, our significant parameters include *µ*_*p*_ as well as the primate latency rate, *ϵ*_*p*_, human movement from the urban cycle rate, *τ*_*u*_, human movement from the peri-urban cycle, *τ*_*p*_, and NHP movement from the sylvatic cycle rate, *φ*_*s*_. A significant difference between total order and first order indices indicates the presence of higher order effects. We see evidence of substantial higher order effects in the urban and peri-urban cycles, indicated by the large discrepancies between the red and blue bars.

For cumulative disease incidence of the hosts in the urban and peri-urban cycles, *ν*_*p*_ and the primate latency rate, *ϵ*_*p*_, had non-trivial total order indices while having zero first order indices. This indicates these parameters have higher order effects only, i.e. they must vary with other parameters to cause significant effects on disease incidence. On the other hand, the rates of human movement from the urban cycle, *τ*_*u*_, human movement from the peri-urban cycle, *τ*_*p*_, and NHP movement from the sylvatic cycle rate, *φ*_*s*_ all had significant first and total order Sobol indices. However, the total order indices for each of these is significantly larger than their first order indices. The urban cycle, as seen in panel (a) of figure 3, has an over 60% difference between the first order and total order indices for *τ*_*u*_ and *τ*_*p*_, while a difference of over 30% for the indices of *φ*_*s*_. The higher order effects of the travel parameters are expected because variation of YFV across the three cycles can only occur with variation across each of the movement parameters, recalling that each simulation begins with YFV only in the sylvatic cycle. Since the epidemic always starts in the sylvatic cycle, movement is required for it to spread to the peri-urban and urban cycles. The higher order effects for *ν*_*p*_ and *µ*_*p*_ on the urban and peri-urban cycles are also expected because they modulate the population of susceptible NHPs in the sylvatic cycle. NHPs who successfully recover from YFV acquire lifelong immunity, so the epidemic can quickly die out in the sylvatic cycle before any movement to the peri-urban or urban cycles, unless there are new recruitment of susceptibles through births.

### 5.2 Two-at-a-time analysis

The first order and total Sobol indices inform us of the presence of first order and higher order effects of a parameter. However, they do not tell us how variation of a parameter quantitatively changes a QoI, e.g. how increasing *ν*_*p*_ increases cumulative disease incidence in the sylvatic cycle. To infer this kind of information, a local analysis is required. As our model is dominated by higher order effects, as evidenced in figure 3, we explore some of them through a local analysis. Recall that the total order indices tell us whether higher order effects are present or not; they do not tell us which higher order effects are present. Thus, we choose to explore a subset of parameter combinations through a two-at-a-time analysis. For this analysis, we fix every parameter but our two of interest. We then vary these two parameters and see how they affect cumulative disease incidence.

For the peri-urban cycle, for both human and NHP disease incidences, we found no discernible relationships from varying the parameters with significant higher order effects two at a time. However, for the urban cycle, we found several clear trends in disease incidence while varying two parameters. We found that varying the travel parameters between the urban and peri-urban cycle caused significant variation of disease incidence in the urban cycle. In figure 4, we see that high levels of disease incidence were associated with low, but non-zero, values of both *τ*_*u*_, travel from the urban cycle to the peri-urban cycle, and *τ*_*p*_, travel from the peri-urban cycle to the urban cycle. Higher incidence is seen with *τ*_*p*_ dominating values of *τ*_*u*_. If *τ*_*u*_ stays low, then *τ*_*p*_, can vary considerably without dramatically changing levels of disease incidence. However, *τ*_*p*_’s dominance fades as *τ*_*u*_ gets larger. For *τ*_*u*_ ≥ 0.25, there is very low disease incidence, regardless the value of *τ*_*p*_. For any *τ*_*p*_ ≤ 0.65, *τ*_*u*_ can be almost zero, and high levels of incidence in the urban cycle still occur. When both *τ*_*u*_ and *τ*_*p*_ are high, the dynamics of both the urban cycle and the human peri-urban compartments become more closely coupled than the dynamics of the human peri-urban and the NHP peri-urban compartments, and so with fixed travel parameters between the sylvatic NHP and the peri-urban NHP, we see no disease incidence in the urban cycle.

**Figure 4:**
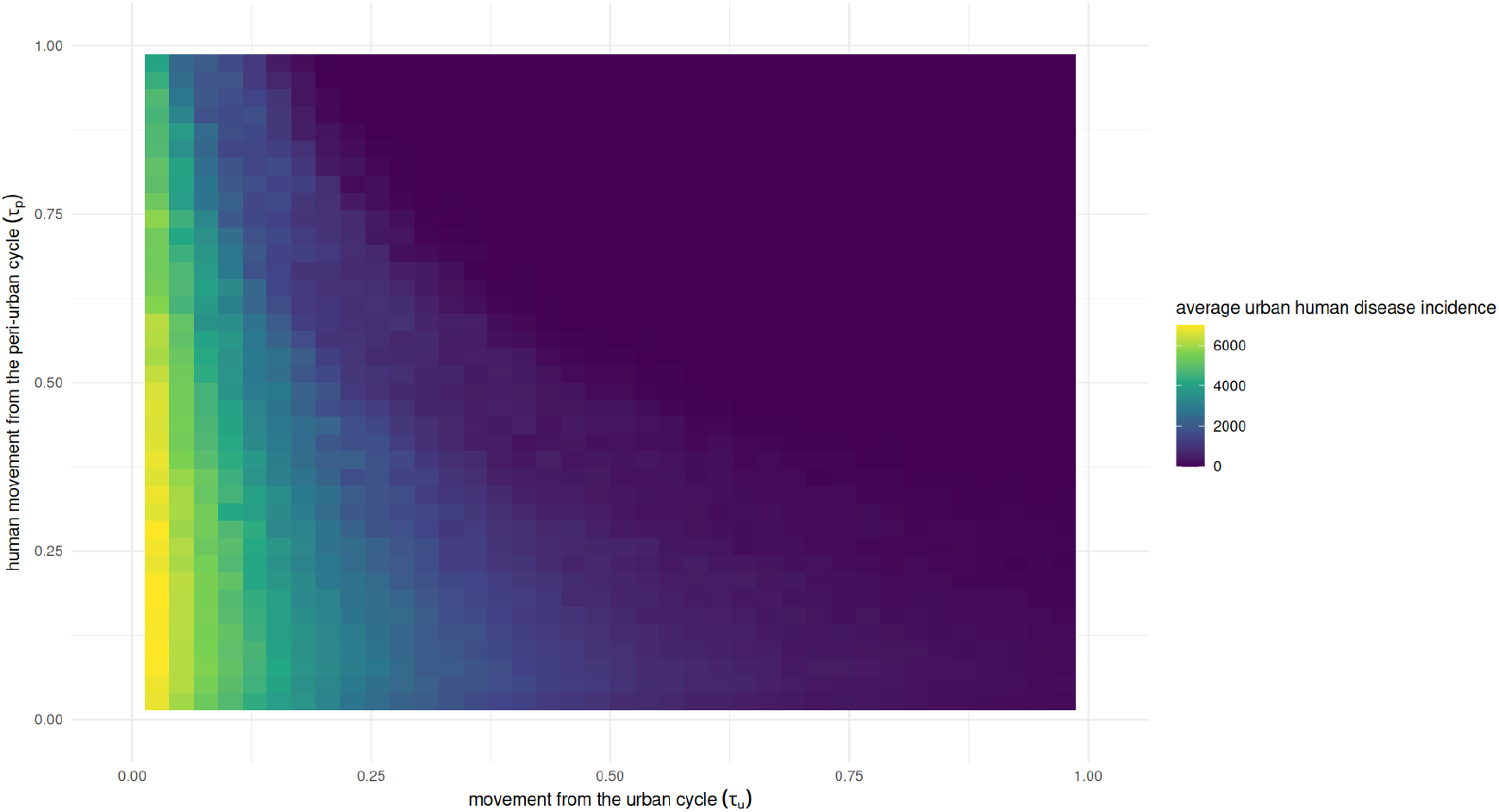
The effects of varying human movement between the urban and periurban cycles on disease incidence in the urban area. If *τ*_*u*_ remains low, *τ*_*p*_ can vary considerably and high levels of disease incidence still occur. However, disease incidence quickly tapers off as *τ*_*u*_ increases, and infections become absent from the urban cycle for high levels of *τ*_*u*_.

We also found an influential relationship between movement parameters and the NHP latency rate. We found that high levels of NHP latency paired with moderate to low levels of movement lead to more incidence in the urban cycle, demonstrated in figure 5. High levels of NHP latency, *ϵ*_*p*_, when coupled with low or moderate levels of movement from the sylvatic cycle, lead to the highest urban incidence rates. The relationship is more tightly coupled for NHP movement from the sylvatic cycle, *φ*_*s*_, than for *τ*_*p*_. Indeed, *ϵ*_*p*_ must be at the tail end of its range, and *φ*_*s*_ roughly between 0.25 and 0.4 for there to be any substantial effect on urban disease incidence. Meanwhile, *τ*_*p*_ can be between 0 and 0.3, and if *ϵ*_*p*_ *>* 0.005, we see significant levels of incidence in the urban cycle.

**Figure 5:**
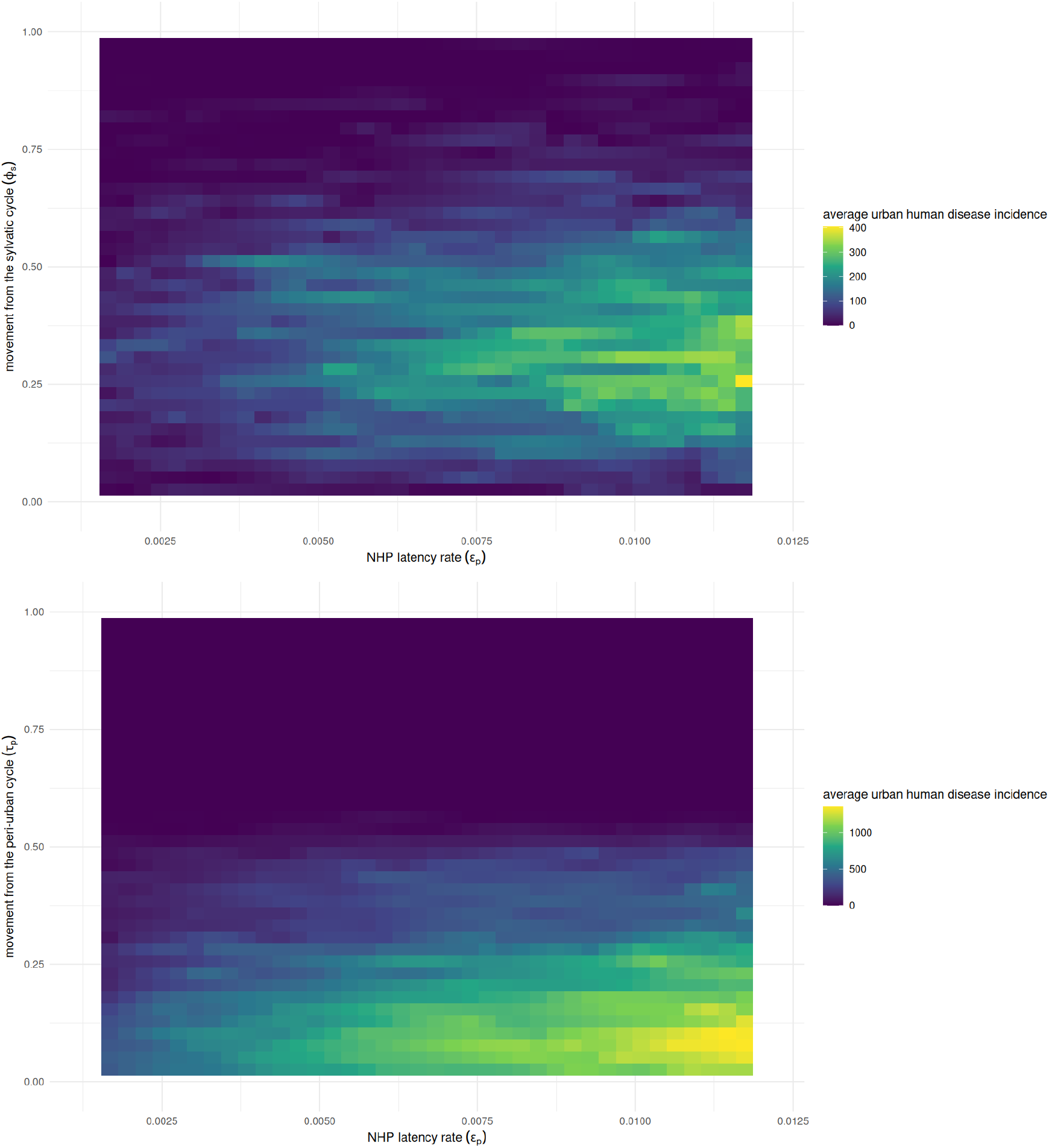
The effects of varying the NHP latency rate with travel rates on urban disease incidence. Low to moderate levels of movement from the peri-urban cycle coupled with short to long periods of latency with moderately high levels of incidence still occurring. The relationship between *φ*_*s*_ and *ϵ*_*p*_ is tighter, with moderately high levels of *ϵ*_*p*_ in conjunction with low to moderate levels of *φ*_*s*_ for relatively high levels of incidence to occur.

## 6 Discussion

The dynamics of emerging infectious diseases are complex and highly stochastic, and modeling this ecology mechanistically requires parameter rich, stochastic models. Often, data on emergence is plagued with large uncertainty, which makes inference on these parameter rich models even more difficult. Understanding how large uncertainties in the parameters propagates to uncertainty in quantities of interest is a key task for the modeler trying to infer mechanisms underlying emergence in a particular disease system. Sobol indices are a statistically rigorous way to quantify the effects of a parameter’s uncertainty on quantities of interest, but their computation has, until recently, been prohibitively expensive for stochastic models. The variance deconvolution estimator allows one to rigorously delineate the variation arising from a stochastic solver apart from the variation arising from the parameter space exploration, allowing for a more accurate and efficient computation of Sobol indices for stochastic models. For the modeler working on emerging infectious diseases, this opens the door for a more robust, mechanistic analysis of emergence. Indeed, without global sensitivity analysis, we are left to either make assumptions about model parameter importance a priori, or we are left to use a local sensitivity analysis, where one varies parameters, or groups of parameters, individually to see their effect on model outcome.

Our YFV model highlights the limitation of these two options. Although our model makes many simplifying assumptions about YFV’s ecology in South America, which we discuss in more detail below, it still contains 28 parameters, making the number of possible parameter combinations impracticable; the number of parameter groupings is 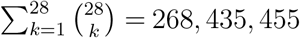. Consequently, one might only perform a one-at-a-time analysis, and for our model, we would have only seen significant variation for the NHP birth and death rate parameters, a modest amount of variation for the travel parameters, and no variation for the NHP latency parameter. However, we know from the difference between the total order indices and the first order indices, i.e. the higher order effects, that uncertainty in our model is heavily characterized by higher order effects, which we would have missed with a local analysis. Thus, computing first order and total order Sobol indices circumvents a combinatorial nightmare while also giving us a global picture of each parameter. If instead we had decided to make assumptions about model parameter importance a priori, we easily could have made erroneous assumptions, even if guided by the underlying biology. We know that in each transmission cycle of YFV, we have a different composition of hosts and vectors. YFV is endemic in the sylvatic cycle, but much less is known about the vector species that help sustain the cycle, which gives us much larger uncertainty in the parameter ranges associated with their life history in our model. We might expect that this uncertainty would have non-trivial effect on disease incidence. However, the Sobol indices demonstrated that this uncertainty was immaterial on the variation of disease incidence across the cycles.

The higher order effects that did have substantial effect on the variation of disease incidence across the cycles, and especially the urban cycle, are largely from interactions between the travel parameters. Recall that these parameters give the linear rate that host individuals move from a compartment in one cycle to another, e.g. *τ*_*u*_*S*_*u*_ is the rate that individuals move from *S*_*u*_ to *S*_*p*_. Since this is a Markov chain, a certain fraction of *S*_*u*_ turns over every 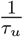 units of time, and we do not expect that host populations actually behave this way. These parameters are phenomenological, making their direct translation difficult. Instead, we interpret these parameters as representing a level of connectivity between the transmission cycles. We explored this connectivity with our two-at-a-time analysis, and we found that low levels of connectivity between the cycles often leads to the high level of cumulative disease incidence in the urban cycle. Here, the peri-urban cycle acts as a highway between the sylvatic cycle and the urban cycle, increasing the likelihood of YFV incidence in urban areas, validating the empirical efforts investigating the ecology of transition zones in YFV transmission [22]. We hypothesize that higher order interactions, beyond second order interactions, are occurring as well. In particular, we hypothesize that variation of disease incidence in the peri-urban cycle is influenced by variation across the travel parameters for human travel, *τ*_*u*_ and *τ*_*p*_, as well as variation across the travel parameters for NHP travel, *φ*_*p*_ and *φ*_*s*_. Efficient algorithms exist to compute higher order indices, and future work will explore these hypotheses by implementing these algorithms in conjunction with the variance deconvolution estimator [50].

The utility of models like ours to explore ecological mechanisms of, or to forecast, YFV transmission and its re-emergence will rely on narrowing the uncertainty of connectivity between the transmission cycles. This raises several important points. First, modeling always makes assumptions, and the inferences drawn from global sensitivity analysis relies on some key assumptions. Sobol indices are computed for particular parameter ranges, and it is entirely possible that exploring a different part of parameter space will lead to different Sobol indices. Furthermore, parameters that have been well studied are more likely to have less uncertainty, and so it is not surprising that a parameter in a narrower range will cause less variation than a parameter with a larger range. A parameter with trivial Sobol indices is not to say it is not an important process ecologically, but rather, that we have narrowed it down to a range that our model and our quantities of interest do not experience substantial variation from. Instead, the Sobol indices suggest where future empirical work would be most useful. When we consider our travel parameters, they have a much larger range compared to other parameters of the model because how the transmission cycles are coupled together is a difficult, open question that requires further attention empirically and mathematically.

This leads us to the second point. In addition to the part of parameter space we choose to explore, the form of our model and our quantities of interest also impact the computation of our Sobol indices. Here, we chose a multiple patch model and cumulative disease incidence to explore how connectivity between the transmission cycles affects the re-emergence of YFV. Yet, other quantities of interest, besides cumulative disease incidence, could have been used to investigate re-emergence. With cumulative disease incidence, we do not look at the size or length of a particular outbreak, so it is possible that some parameters’ uncertainties are far more consequential on the number of outbreaks and the duration of each outbreak than on cumulative disease incidence during the simulation.

The threat of emerging infectious diseases increasingly requires more realistic models to explore mitigation and control efforts as well as forecast future scenarios. This has led the modeling community towards more complex models that are often stochastic. Until recently, global sensitivity analysis was infeasible for this class of models, due simply to computational constraints. Here, we demonstrated how recent developments from uncertainty quantification have allowed us to circumvent these constraints. Providing this rigorous statistical analysis of our models, these tools will continue to play an increasingly important role in modeling emerging infectious diseases.

## 7 Acknowledgments

We would like to thank the UNM Center for Advanced Research Computing, supported in part by the National Science Foundation, for providing the high performance computing resources used in this work. We thank Owen Lewis for discussions about global sensitivity anlysis, and we also thank Owen Davis and Teresa Portone for suggesting the use of the variance deconvolution for our sensitivity analysis.

## References

1. Anderson RM and May RM. Infectious Diseases of Humans: Dynamics and Control. Oxford University Press, 1991 May 16. doi: 10.1093/oso/9780198545996.001.0001. Available from: https://doi.org/10.1093/oso/9780198545996.001.0001 [Accessed on: 2026 Jan 21]

2. Keeling MJ and Rohani P. Modeling Infectious Diseases in Humans and Animals — Princeton University Press. ISBN: 9780691116174. 2007 Oct 28. Available from: https://press.princeton.edu/books/hardcover/9780691116174/modeling-infectious-diseases-in-humans-and-animals [Accessed on: 2025 Aug 18]

3. Brett TS and Rohani P. Transmission dynamics reveal the impracticality of COVID-19 herd immunity strategies. Proceedings of the National Academy of Sciences. 2020 Oct 13; 117. Publisher: Proceedings of the National Academy of Sciences:25897–903. doi: 10.1073/pnas.2008087117. Available from: https://www.pnas.org/doi/abs/10.1073/pnas.2008087117 [Accessed on: 2025 Nov 4]

4. Gokhale DV, Brett TS, He B, King AA, and Rohani P. Disentangling the causes of mumps reemergence in the United States. Proceedings of the National Academy of Sciences. 2023 Jan 17; 120. Publisher: Proceedings of the National Academy of Sciences:e2207595120. doi: 10.1073/pnas.2207595120. Available from: https://www.pnas.org/doi/abs/10.1073/pnas.2207595120 [Accessed on: 2025 Nov 4]

5. Allen LJS. A primer on stochastic epidemic models: Formulation, numerical simulation, and analysis. Infectious Disease Modelling. 2017 May 1; 2:128–42. doi: 10.1016/j.idm.2017.03.001. Available from: https://www.sciencedirect.com/science/article/pii/S2468042716300495 [Accessed on: 2025 Aug 18]

6. King AA, Domenech de Cellès M, Magpantay FMG, and Rohani P. Avoidable errors in the modelling of outbreaks of emerging pathogens, with special reference to Ebola. Proceedings of the Royal Society B: Biological Sciences. 2015 May 7; 282. Publisher: Royal Society:20150347. doi: 10.1098/rspb.2015.0347. Available from: https://royalsocietypublishing.org/doi/10.1098/rspb.2015.0347 [Accessed on: 2025 Aug 18]

7. Gillespie DT. Exact stochastic simulation of coupled chemical reactions. The Journal of Physical Chemistry. 1977 Dec 1; 81. Publisher: American Chemical Society:2340–61. doi: 10.1021/j100540a008. Available from: https://doi.org/10.1021/j100540a008 [Accessed on: 2025 Aug 18]

8. Saltelli A, Ratto M, Andres T, Campolongo F, Cariboni J, Gatelli D, Saisana M, and Tarantola S. Global Sensitivity Analysis. The Primer — Wiley Online Books. 2007 Dec 18. Available from: https://onlinelibrary.wiley.com/doi/book/10.1002/9780470725184 [Accessed on: 2025 Aug 18]

9. Hart JL, Alexanderian A, and Gremaud PA. Efficient Computation of Sobol’ Indices for Stochastic Models. SIAM Journal on Scientific Computing. 2017 Jan; 39. Publisher: Society for Industrial and Applied Mathematics:A1514–A1530. doi: 10.1137/16M106193X. Available from: https://epubs.siam.org/doi/10.1137/16M106193X [Accessed on: 2025 Aug 18]

10. Zhu X and Sudret B. Global sensitivity analysis for stochastic simulators based on generalized lambda surrogate models. Reliability Engineering & System Safety. 2021 Oct 1; 214:107815. doi: 10.1016/j.ress.2021.107815. Available from: https://www.sciencedirect.com/science/article/pii/S0951832021003379 [Accessed on: 2025 Aug 18]

11. Clements KB, Geraci G, Olson AJ, and Palmer TS. A variance deconvolution estimator for efficient uncertainty quantification in Monte Carlo radiation transport applications. Journal of Quantitative Spectroscopy and Radiative Transfer. 2024 Jun 1; 319:108958. doi: 10.1016/j.jqsrt.2024.108958. Available from: https://www.sciencedirect.com/science/article/pii/S0022407324000657 [Accessed on: 2025 Aug 18]

12. Clements K, Geraci G, Olson AJ, and Palmer TS. Global sensitivity analysis in Monte Carlo radiation transport. 2024 Mar 16. doi: 10.48550/arXiv.2403.06106. arXiv: 2403.06106[physics]. Available from: http://arxiv.org/abs/2403.06106 [Accessed on: 2025 Aug 18]

13. Chippaux JP and Chippaux A. Yellow fever in Africa and the Americas: a historical and epidemiological perspective. The Journal of Venomous Animals and Toxins Including Tropical Diseases. 2018; 24:20. doi: 10.1186/s40409-018-0162-y

14. Gianchecchi E, Cianchi V, Torelli A, and Montomoli E. Yellow Fever: Origin, Epidemiology, Preventive Strategies and Future Prospects. Vaccines. 2022 Feb 27; 10:372. doi: 10.3390/vaccines10030372

15. Oliveira Figueiredo P de, Stoffella-Dutra AG, Barbosa Costa G, Silva de Oliveira J, Dourado Amaral C, Duarte Santos J, Soares Rocha KL, Araújo Júnior JP, Lacerda Nogueira M, Zazá Borges MA, Pereira Paglia A, Desiree LaBeaud A, Santos Abrahão J, Geessien Kroon E, Bretas de Oliveira D, Paiva Drumond B, and Souza Trindade G de. Re-Emergence of Yellow Fever in Brazil during 2016–2019: Challenges, Lessons Learned, and Perspectives. Viruses. 2020 Nov; 12. Publisher: Multidisciplinary Digital Publishing Institute:1233. doi: 10.3390/v12111233. Available from: https://www.mdpi.com/1999-4915/12/11/1233 [Accessed on: 2025 Aug 18]

16. Sacchetto L, Drumond BP, Han BA, Nogueira ML, and Vasilakis N. Re-emergence of yellow fever in the neotropics - quo vadis? Emerging Topics in Life Sciences. 2020 Dec 11; 4:399–410. doi: 10.1042/ETLS20200187

17. Cunha MdP, Duarte-Neto AN, Pour SZ, Ortiz-Baez AS, Černý J, Pereira BBdS, Braconi CT, Ho YL, Perondi B, Sztajnbok J, Alves VAF, Dolhnikoff M, Holmes EC, Saldiva PHN, and Zanotto PMdA. Origin of the São Paulo Yellow Fever epidemic of 2017–2018 revealed through molecular epidemiological analysis of fatal cases. Scientific Reports. 2019 Dec 31; 9. Publisher: Nature Publishing Group:20418. doi: 10.1038/s41598-019-56650-1. Available from: https://www.nature.com/articles/s41598-019-56650-1 [Accessed on: 2025 Aug 18]

18. Bryant JE, Holmes EC, and Barrett ADT. Out of Africa: A Molecular Perspective on the Introduction of Yellow Fever Virus into the Americas. PLOS Pathogens. 2007 May 18; 3. Publisher: Public Library of Science:e75. doi: 10.1371/journal.ppat.0030075. Available from: https://journals.plos.org/plospathogens/article?id=10.1371/journal.ppat.0030075 [Accessed on: 2025 Aug 18]

19. Despommier D, Ellis BR, and Wilcox BA. The Role of Ecotones in Emerging Infectious Diseases. Ecohealth. 2006; 3:281–9. doi: 10.1007/s10393-006-0063-3. Available from: https://www.ncbi.nlm.nih.gov/pmc/articles/PMC7088109/ [Accessed on: 2025 Aug 18]

20. Borremans B, Faust C, Manlove KR, Sokolow SH, and Lloyd-Smith JO. Cross-species pathogen spillover across ecosystem boundaries: mechanisms and theory. Philosophical Transactions of the Royal Society of London Series B, Biological Sciences. 2019 Sep 30; 374:20180344. doi: 10.1098/rstb.2018.0344

21. Hendy A, Hernandez-Acosta E, Chaves BA, Fé NF, Valério D, Mendonça C, Lacerda MVGd, Buenemann M, Vasilakis N, and Hanley KA. Into the woods: Changes in mosquito community composition and presence of key vectors at increasing distances from the urban edge in urban forest parks in Manaus, Brazil. Acta Tropica. 2020 Jun 1; 206:105441. doi: 10.1016/j.actatropica.2020.105441. Available from: https://www.sciencedirect.com/science/article/pii/S0001706X20300206 [Accessed on: 2025 Aug 18]

22. Hendy A, Hernandez-Acosta E, Valério D, Fé NF, Mendonça CR, Costa ER, Andrade ESd, Júnior JTA, Assunção FP, Scarpassa VM, Lacerda MVGd, Buenemann M, Vasilakis N, and Hanley KA. Where boundaries become bridges: Mosquito community composition, key vectors, and environmental associations at forest edges in the central Brazilian Amazon. PLOS Neglected Tropical Diseases. 2023 Apr 26; 17. Publisher: Public Library of Science:e0011296. doi: 10.1371/journal.pntd.0011296. Available from: https://journals.plos.org/plosntds/article?id=10.1371/journal.pntd.0011296 [Accessed on: 2025 Aug 18]

23. Garcia-Oliveira GF, Guimarães ACDS, Moreira GD, Costa TA, Arruda MS, Mello ÉM de, Silva MC, Almeida MG de, Hanley KA, Vasilakis N, and Drumond BP. YELLOW ALERT: Persistent Yellow Fever Virus Circulation among Non-Human Primates in Urban Areas of Minas Gerais State, Brazil (2021–2023). Viruses. 2024 Jan; 16. Publisher: Multidisciplinary Digital Publishing Institute:31. doi: 10.3390/v16010031. Available from: https://www.mdpi.com/1999-4915/16/1/31 [Accessed on: 2025 Aug 18]

24. Althouse BM, Lessler J, Sall AA, Diallo M, Hanley KA, Watts DM, Weaver SC, and Cummings DAT. Synchrony of sylvatic dengue isolations: a multi-host, multi-vector SIR model of dengue virus transmission in Senegal. PLoS neglected tropical diseases. 2012; 6:e1928. doi: 10.1371/journal.pntd.0001928

25. Althouse BM, Vasilakis N, Sall AA, Diallo M, Weaver SC, and Hanley KA. Potential for Zika Virus to Establish a Sylvatic Transmission Cycle in the Americas. PLoS Neglected Tropical Diseases. 2016 Dec 15; 10:e0005055. doi: 10.1371/journal.pntd.0005055. Available from: https://www.ncbi.nlm.nih.gov/pmc/articles/PMC5157942/ [Accessed on: 2025 Aug 18]

26. Childs ML, Nova N, Colvin J, and Mordecai EA. Mosquito and primate ecology predict human risk of yellow fever virus spillover in Brazil. Philosophical Transactions of the Royal Society of London Series B, Biological Sciences. 2019 Sep 30; 374:20180335. doi: 10.1098/rstb.2018.0335

27. Smith DL, Battle KE, Hay SI, Barker CM, Scott TW, and McKenzie FE. Ross, Macdonald, and a Theory for the Dynamics and Control of Mosquito-Transmitted Pathogens. PLOS Pathogens. 2012 Apr 5; 8. Publisher: Public Library of Science:e1002588. doi: 10.1371/journal.ppat.1002588. Available from: https://journals.plos.org/plospathogens/article?id=10.1371/journal.ppat.1002588 [Accessed on: 2025 Aug 18]

28. Fernandes NCdA, Guerra JM, Díaz-Delgado J, Cunha MS, Saad Ld, Iglezias SD, Ressio RA, Cirqueira CdS, Kanamura CT, Jesus IP, Maeda AY, Vasami FG, Carvalho Jd, Araújo LJd, Souza RPd, Nogueira JS, Spinola RM, and Catão-Dias JL. Differential Yellow Fever Susceptibility in New World Nonhuman Primates, Comparison with Humans, and Implications for Surveillance. Emerging Infectious Diseases. 2021 Jan; 27:47. doi: 10.3201/eid2701.191220. Available from: https://pmc.ncbi.nlm.nih.gov/articles/PMC7774563/ [Accessed on: 2025 Aug 18]

29. Gillespie DT. Stochastic simulation of chemical kinetics. Annual Review of Physical Chemistry. 2007; 58:35–55. doi: 10.1146/annurev.physchem.58.032806.104637

30. Tian T and Burrage K. Binomial leap methods for simulating stochastic chemical kinetics. The Journal of Chemical Physics. 2004 Dec 1; 121:10356–64. doi: 10.1063/1.1810475. Available from: https://doi.org/10.1063/1.1810475 [Accessed on: 2025 Aug 18]

31. Andrade AC de. Density of marmosets in highly urbanised areas and the positive effect of arboreous vegetation. Urban Ecosystems. 2022 Feb 1; 25:101–9. doi: 10.1007/s11252-021-01131-5. Available from: https://doi.org/10.1007/s11252-021-01131-5 [Accessed on: 2025 Aug 18]

32. Hue T, Caubet M, and Moura ACdA. Howlers and marmosets in Pacatuba: an over-crowded existence in a semi-deciduous Atlantic forest fragment? Mammalia. 2017 Jul 1; 81. Publisher: De Gruyter:339–48. doi: 10.1515/mammalia-2015-0167. Available from: https://www.degruyterbrill.com/document/doi/10.1515/mammalia-2015-0167/html?lang=en&srsltid=AfmBOoqhSmJv05ZMJ2KWh4YWJSSn5q_AhUJ9czTOY6e0Wct5M20X0-8y [Accessed on: 2025 Aug 18]

33. Brazil BR: Rural Population — Economic Indicators — CEIC. Available from: https://www.ceicdata.com/en/brazil/population-and-urbanization-statistics/br-rural-population [Accessed on: 2025 Aug 18]

34. São Paulo (Municipality, Brazil)- Population Statistics, Charts, Map and Location. Available from: https://www.citypopulation.de/en/brazil/regiaosudeste/admin/s%C3%A3o_paulo/3550308s%C3%A3o_paulo/ [Accessed on: 2025 Aug 18]

35. Ravetz J and Sahana M. Chapter 1 - Where is the peri-urban? A working definition and global typology. Modern Cartography Series. Ed. by Sahana M. Vol. 11. Remote Sensing and GIS in Peri-Urban Research. Academic Press, 2024 Jan 1:3–25. doi: 10.1016/B978-0-443-15832-2.00001-0. Available from: https://www.sciencedirect.com/science/article/pii/B9780443158322000010 [Accessed on: 2025 Aug 18]

36. Sobol IM. Global sensitivity indices for nonlinear mathematical models and their Monte Carlo estimates. Mathematics and Computers in Simulation. The Second IMACS Seminar on Monte Carlo Methods 2001 Feb 15; 55:271–80. doi: 10.1016/S0378-4754(00)00270-6. Available from: https://www.sciencedirect.com/science/article/pii/S0378475400002706 [Accessed on: 2025 Aug 18]

37. Alouatta caraya - Vertebrate Collection — UWSP. Available from: https://www3.uwsp.edu:443/biology/VertebrateCollection/Pages/Vertebrates/Mammals%20of%20Paraguay/Alouatta%20caraya/Alouatta%20caraya.aspx [Accessed on: 2025 Aug 18]

38. Common marmoset. Wisconsin National Primate Research Center. Available from: https://primate.wisc.edu/primate-info-net/pin-factsheets/pin-factsheet-common-marmoset/ [Accessed on: 2025 Aug 18]

39. Rumiz DI. Alouatta caraya: Population density and demography in Northern Argentina. American Journal of Primatology. 1990; 21:279–94. doi: 10.1002/ajp.1350210404

40. Common marmoset. Wisconsin National Primate Research Center. Available from: https://primate.wisc.edu/primate-info-net/pin-factsheets/pin-factsheet-common-marmoset/ [Accessed on: 2025 Aug 18]

41. Sowilem MM, Kamal HA, and Khater EI. Life table characteristics of Aedes aegypti (Diptera:Culicidae) from Saudi Arabia. Tropical Biomedicine. 2013 Jun; 30:301–14

42. Nur Aida H, Abu Hassan A, Nurita AT, Che Salmah MR, and Norasmah B. Population analysis of Aedes albopictus (Skuse) (Diptera:Culicidae) under uncontrolled laboratory conditions. Tropical Biomedicine. 2008 Aug; 25:117–25

43. Captain-Esoah M, Kweku Baidoo P, Frempong KK, Adabie-Gomez D, Chabi J, Obuobi D, Kwame Amlalo G, Balungnaa Veriegh F, Donkor M, Asoala V, Behene E, Adjei Boakye D, and Dadzie SK. Biting Behavior and Molecular Identification of Aedes aegypti (Diptera: Culicidae) Subspecies in Some Selected Recent Yellow Fever Outbreak Communities in Northern Ghana. Journal of Medical Entomology. 2020 Jul 4; 57:1239–45. doi: 10.1093/jme/tjaa024

44. Liu H, Liu L, Cheng P, Yang L, Chen J, Lu Y, Wang H, Chen XG, and Gong M. Bionomics and insecticide resistance of Aedes albopictus in Shandong, a high latitude and high-risk dengue transmission area in China. Parasites & Vectors. 2020 Jan 9; 13:11. doi: 10.1186/s13071-020-3880-2. Available from: https://doi.org/10.1186/s13071-020-3880-2 [Accessed on: 2025 Aug 18]

45. Stanzani LMdA, Motta MdA, Erbisti RS, Abreu FVSd, Nascimento-Pereira AC, Ferreira-de-Brito A, Neves MSAS, Pereira GR, Pereira GR, Santos CBD, Pinto IdS, Vicente CR, Faccini-Martínez ÁA, Cavalcante KRLJ, Falqueto A, and Lourenço-de-Oliveira R. Back to Where It Was First Described: Vectors of Sylvatic Yellow Fever Transmission in the 2017 Outbreak in Espírito Santo, Brazil. Viruses. 2022 Dec 15; 14:2805. doi: 10.3390/v14122805

46. Couto-Lima D, Madec Y, Bersot MI, Campos SS, Motta MdA, Santos FBD, Vazeille M, Vasconcelos PFdC, Lourenço-de-Oliveira R, and Failloux AB. Potential risk of re-emergence of urban transmission of Yellow Fever virus in Brazil facilitated by competent Aedes populations. Scientific Reports. 2017 Jul 7; 7:4848. doi: 10.1038/s41598-017-05186-3

47. CDC. Yellow Fever. Yellow Book. 2025 Jun 23. Available from: https://www.cdc.gov/yellow-book/hcp/travel-associated-infections-diseases/yellow-fever.html [Accessed on: 2025 Aug 18]

48. Silva NIO, Sacchetto L, Rezende IM de, Trindade GdS, LaBeaud AD, Thoisy B de, and Drumond BP. Recent sylvatic yellow fever virus transmission in Brazil: the news from an old disease. Virology Journal. 2020 Jan 23; 17:9. doi: 10.1186/s12985-019-1277-7

49. Yellow Fever - PAHO/WHO — Pan American Health Organization. 2025 Aug 14. Available from: https://www.paho.org/en/topics/yellow-fever [Accessed on: 2025 Aug 18]

50. Saltelli A. Making best use of model evaluations to compute sensitivity indices. Computer Physics Communications. 2002 May 15; 145:280–97. doi: 10.1016/S0010-4655(02)00280-1. Available from: https://www.sciencedirect.com/science/article/pii/S0010465502002801 [Accessed on: 2025 Aug 18]

